# More Reliable EEG Electrode Digitizing Methods Can Reduce Source Estimation Uncertainty, But Current Methods Already Accurately Identify Brodmann Areas

**DOI:** 10.1101/557074

**Authors:** Seyed Yahya Shirazi, Helen J. Huang

**Author notes:** S.Y. Shirazi, (corresponding author, email). H.J. Huang (email:).

## Abstract

Electroencephalography (EEG) and source estimation can be used to identify brain areas activated during a task, which could offer greater insight on cortical dynamics. Source estimation requires knowledge of the locations of the EEG electrodes. This could be provided with a template or obtained by digitizing the EEG electrode locations. Operator skill and inherent uncertainties of a digitizing system likely produce a range of digitization reliabilities, which could affect source estimation and the interpretation of the estimated source locations. Here, we compared the reliability of five digitizing methods (ultrasound, structured-light 3D scan, infrared 3D scan, motion capture probe, and motion capture) and determined the relationship between digitization reliability and source estimation uncertainty, assuming other contributors to source estimation uncertainty were constant. We digitized a mannequin head using each method five times and quantified the reliability and validity of each method. We created five hundred sets of electrode locations based on our reliability results and applied a dipole fitting algorithm (DIPFIT) to perform source estimation. The motion capture method, which recorded the locations of markers placed directly on the electrodes had the best reliability with an average electrode variability of 0.001cm. Then, in order of decreasing reliability were the method using a digitizing probe in the motion capture system, an infrared 3D scanner, a structured-light 3D scanner, and an ultrasound digitization system. Unsurprisingly, uncertainty of the estimated source locations increased with greater variability of EEG electrode locations and less reliable digitizing systems. If EEG electrode location variability was ~ 1 cm, a single source could shift by as much as 2 cm. To help translate these distances into practical terms, we quantified Brodmann area accuracy for each digitizing method and found that the average Brodmann area accuracy for all digitizing methods was > 80%. Using a template of electrode locations reduced the Brodmann area accuracy to ~ 50%. Overall, more reliable digitizing methods can reduce source estimation uncertainty, but the significance of the source estimation uncertainty depends on the desired spatial resolution. For accurate Brodmann area identification, any of the digitizing methods tested can be used confidently.

## I. INTRODUCTION

Estimating active cortical sources using electroencephalography (EEG) is becoming widely adopted in multiple research areas as a non-invasive and mobile functional brain imaging modality [1]–[4]. EEG is the recording of the electrical activity on the scalp and is appealing for studying cortical dynamics during movements and decision making due to the high temporal (i.e. millisecond) resolution of electrical signals. One of the challenges of using EEG is that the signal recorded in an EEG electrode is a mixture of electrical activity from multiple sources, which include the cortex, muscles, heart, eye, 60 Hz noise from power lines, and motion artifacts from cable sway and head movements [5], [6]. To meaningfully correlate EEG analyses with brain function, the unwanted source content such as muscle activity, eye blinks, and motion artifacts need to be attenuated or separated from the cortical signal content. A multitude of tools such as independent component analysis, artifact rejection algorithms, and phantom heads have been developed to address the need to separate the source signals to extract the underlying cortical signal [7]–[11]. Using high-density EEG and improving EEG post-processing techniques have also improved spatial resolution of source estimation to ~ 1 cm in experimental studies [12]–[16].

Source estimation requires knowing the EEG signals and the locations of the EEG electrodes to estimate the locations of the cortical sources that produced the EEG signals measured on the scalp. An intuitive assumption of source estimation is that precise placement of the EEG electrodes on the scalp is essential for accurate estimation of source locations [17]. Computational studies reported shifts of 0.5 cm to 1.2 cm in estimated source locations as a result of 0.5 cm (or 5^°^) error in the electrode digitization [18]–[22]. For EEG studies conducted inside a magnetic resonance imaging (MRI) device, the exact electrode locations with respect to the cortex can be captured and processed, which results in near perfect alignment of identified brain areas [14], [23]. However, for studies that do not involve MRI, the electrode locations should be “digitized”, i.e. recorded digitally via a three-dimensional (3D) position recording method [24]. These digitized locations can then be coupled with either a subject-specific or an averaged template of the brain structure obtained from MRI or other imaging techniques to perform EEG source estimation.

Just one decade ago in the mid-2000’s, the main digitizing technologies available were based on ultrasound and electromagnetism, which were expensive, time consuming, and needed trained operators [24], [25]. An ultrasound digitizing system uses differences in ultrasound-wave travel times from emitters on a digitizing wand to an array of receivers to estimate the 3D location of the emitters and the tip of the digitizing wand. An electromagnetic system tracks the locations of receivers on a wand in an emitted electromagnetic field to estimate the position of the tip of the wand. The environment must be clear of magnetic objects when using an electromagnetic digitizing system, otherwise the electrode locations will be warped [26], [27].

Recent efforts have focused on developing technologies to make digitization more accessible and convenient, mainly by incorporating image-based technologies [28], [29]. For example, using photogrammetry and motion capture methods for digitization can provide accurate electrode locations in a short period of time [30], [31]. Photogrammetry involves using cameras to take a series of color images at different view angles. These images can then be analyzed to identify the locations of specific points in the 3D space [30], [32]. Motion capture typically uses multiple infrared cameras around the capture volume to take simultaneous images to identify the locations of reflective or emitting markers. If markers are placed directly on the EEG electrodes, a motion capture system could conveniently record the position of all of the electrodes at once [26], [31]. Motion capture could also be used to record the position of the tip of a probe, a rigid body with multiple markers, to digitize 3D locations of any point in the capture volume. Several recent commercial digitizing systems use simple motion capture approaches to digitize EEG electrode locations with or without a probe [27], [33]–[35].

Another option for digitizing EEG electrodes that has also gained much interest recently are 3D scanners. A common approach for 3D scanning is detecting the infrared or visible reflections of projected light patterns with a camera to estimate the shape of an object [36]. The 3D scanned shapes can then be plotted in a software program such as MATLAB, and the locations of specific points on the 3D scanned shape can be determined. Recently, common EEG analysis toolboxes such as EEGLAB [37] and FieldTrip [38] support using 3D scanners to digitize the electrode locations. Studies suggest that 3D scanners can improve digitization accuracy and significantly reduce digitization time [39]. Using other camera-based systems such as time-of-flight scanners and virtual reality headsets were also reported to provide comparable digitization reliability as the ultrasound or electromagnetic digitizing methods, while reducing the time spent for digitizing the EEG electrodes [27], [40], [41].

The purposes of this study were 1) to compare the reliability and validity of five digitizing methods and 2) to quantify the relationship between digitization reliability and source estimation uncertainty. We determined source estimation uncertainty using spatial metrics and Brodmann areas. We hypothesized that digitizing methods with less reliability would increase uncertainty in the estimates of the electrocortical source locations. For our analyses, we assumed that all other contributors to source estimation uncertainty such as variability of head-meshes and assumptions of electrical conductivity values were constant.

## II. METHODS

We fitted a mannequin head with a 128-channel EEG cap (ActiveTwo EEG system, BioSemi B.V., Amsterdam, the Netherlands, Figure 1A) and used this mannequin head setup to record multiple digitizations of the locations of the EEG electrodes and fiducials (right preauricular, left preauricular, and nasion). To prevent the cap from moving from digitization to digitization, we taped the cap to the mannequin head (Figure 1A). To help ensure that the fiducials were digitized at the same locations for every digitizing method, we marked the fiducials with small 4-mm markers on the mannequin head and with small o-rings on the cap (Figure 1A).

**Fig. 1.**
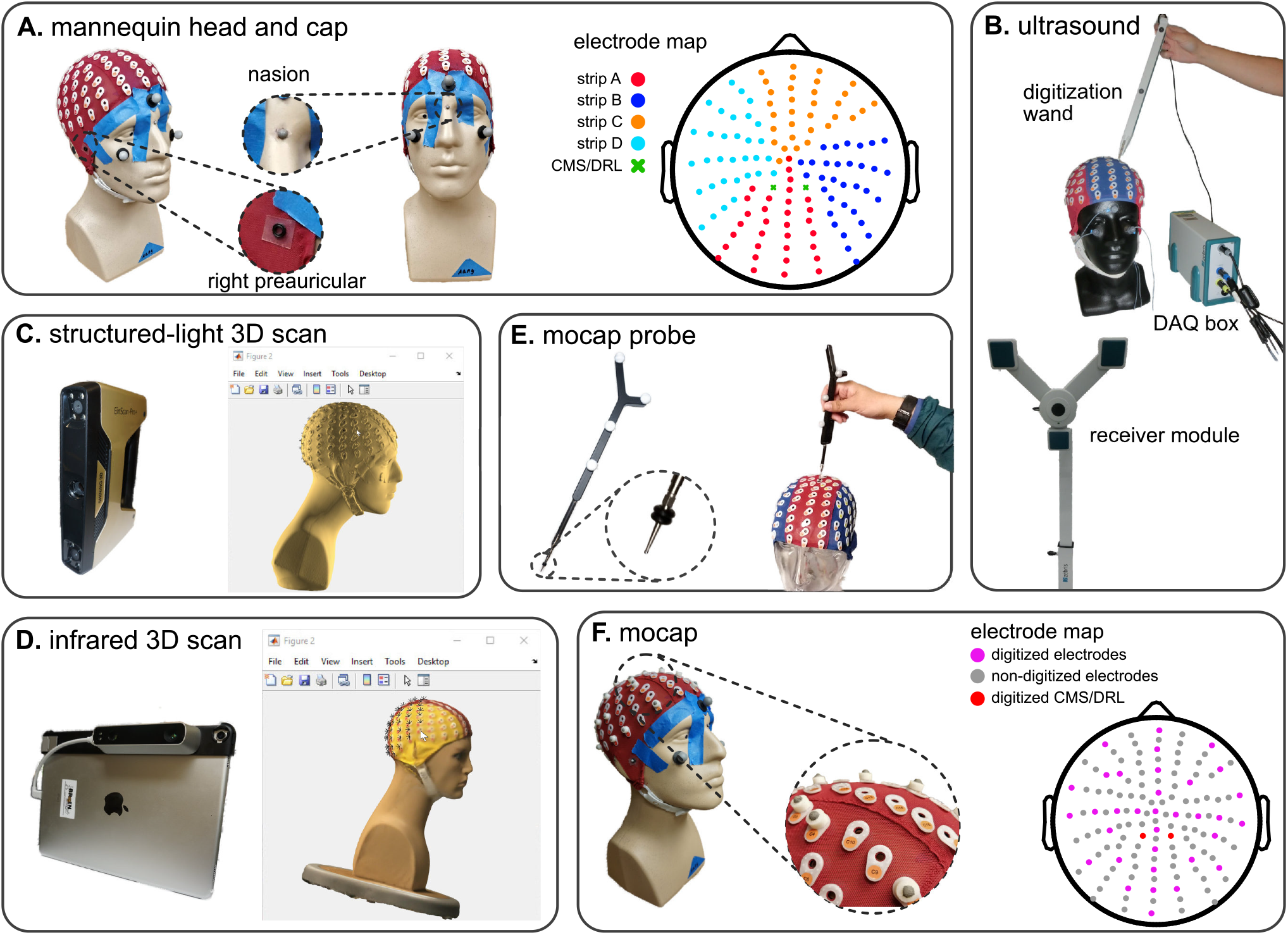
The mannequin head used for digitization and the five digitizing methods tested. **A**: The mannequin head fit with the 128-electrode EEG cap (ActiveTwo EEG system, BioSemi B.V., Amsterdam, the Netherlands) that was used for all of the digitizing recordings. Reflective markers were placed on the face and fiducials (naison, left & right preauricular), which were visible during the digitizing process. The color-coded map of the cap shows the different electrode strips and the order of digitization from A to D. **B**: The ultrasound digitizing system, Zebris, and an operator placing the tip of the wand in the electrode well on the cap. **C**: The structured-light 3D scanner, the EinScan Pro+ and an operator manually marking the locations of individual electrodes of the 3D scan of the mannequin head from the EinScan Pro+ in MATLAB. **D**: The infrared 3D scanner, the Structure Sensor integrated with an Apple^®^ iPad Pro and an operator manually marking the locations of individual electrodes of the colored 3D scan of the mannequin head from the Structure Sensor in MATLAB. **E**: The motion capture digitizing probe with a close-up view of the o-rings placed 7 millimeters away from the tip. The probe functions similarly to the wand in the Zebris ultrasound system. **F**: The EEG cap with 35 3D-printed EEG electrode shaped reflective markers, 3 face markers, and 3 fiducial markers used for the motion capture digitization. The electrode map depicts the approximate locations of the digitized electrodes and grounds (BioSemi CMS and DRL electrodes) using the motion capture method.

### A. Digitizing methods

We compared five methods for digitization: ultrasound, structured-light 3D scanning, infrared 3D scanning, motion capture with a digitizing probe, and motion capture with reflective markers.

For each method, members of the laboratory digitized the mannequin head five times, completing all digitizations in a single session. All of the operators had prior experience in digitizing and were asked to follow each method’s specific guidelines. We imported the digitization data to MATLAB (version 9.4, R2018a, Mathworks, Natick, MA) and performed all analyses in MATLAB.

#### 1) Ultrasound

We used a Zebris positioning system with ElGuide software version 1.6 (Zebris Medical GmBH, Tübingen, Germany, Figure 1B) to digitize the electrodes with an ultrasound method. Following the Zebris manual, we placed 3 ultrasound emitters on the face of the mannequin head, placed the receiver module in front of the mannequin head, and used the digitizing wand to record the electrode locations. We calibrated the system using the ElGuide calibration procedure. We marked the fiducials repeatedly until we obtained fiducials with a digitized 3D location of nasion that was < 2mm with respect to the midline and with preauriculars that had a difference of < 5mm in the anterior/posterior and top/bottom directions. Operators followed the interactive ElGuide template to digitize each electrode location. This process involved fully placing the wand tip into the electrode wells and ensuring that the receivers were able to see all emitters at the time of recording electrode locations, so that the estimated position of the wand tip was stable.

#### 2) Structured-light 3D scan

We used an Einscan Pro+ (Shining 3D Tech. Co. Ltd., Hangzhou, China, Figure 1C) to digitize the electrodes with a structured-light 3D scanner. This scanner estimates the shape of an object from reflections of the projected visible lights. We calibrated the Einscan Pro+ one time with the Einscan’s calibration board and followed the software’s step-by-step instructions. We used the scanner’s hand-held rapid mode with high details and allowed the scanner to track both texture and markers during the scanning process. Each operator scanned the mannequin head until the scan included the cap, fiducials, and the face. We then applied the watertight model option to the scan and exported the model as a `PLY` file to continue the digitization process in MATLAB.

After acquiring the 3D scan, the 3D head model needed to be imported into a software program, where the operator manually marked the EEG electrode locations on the 3D scanned head model. We followed the FieldTrip toolbox documentation for digitization using 3D scanners [42] and created a MATLAB script file for importing and digitizing 3D models of the mannequin head. The operator first marked the fiducials on the mannequin head model in MATLAB to build up the head coordinate system. Then, the operator marked the locations of the electrodes on the screen in each section of the cap in alphanumerical order (A, B, C, D, and the fiducials, total: 131 locations, Figure 1A). The operators referred to a physical EEG cap for guidance to help mark the locations in the expected order because these scans were not in color and the letter labels of the electrodes were not visible on the 3D model.

#### 3) Infrared 3D scan

We used the Structure sensor (model ST01, Occipital Inc., San Francisco, CA) integrated with an Apple^®^ iPad (10-inch Pro) to digitize the electrodes with an infrared dot-projection 3D scanner (Figure 1D). This scanner shares similar working principles as a structured-light scanner but uses infrared light projection to estimate the shape of objects. We calibrated the sensor in daylight and office light according to the manual. We scanned the head using the high color and mesh resolutions. When the mannequin head was completely in the sensor’s field of view, the operator started scanning. The Structure sensor interface gives the operator visual feedback to help the operator obtain a complete high-quality scan. We visually inspected that the scanned model matched the mannequin and then exported the model to the MATLAB environment.

We used the FieldTrip toolbox to import and digitize the 3D mannequin head scans following the same procedure described for the structured-light 3D scan digitization.

#### 4) Motion capture probe

We used a digitizing probe and a 22-camera motion capture system (OptiTrack, Corvallis, OR) to digitize the electrodes. The probe is a solid rigid body with four fixed reflective markers (Figure 1E). We placed three reflective makers on the face of the mannequin to account for possible movements of the head during data collection. Each operator digitized the fiducials and each section of the cap (A, B, C, D, Figure 1A) in separate takes. We placed double o-rings seven millimeters away from the probe tip to ensure consistent placement of the tip inside the electrode wells (Figure 1E). The tracking error of the motion capture system was less than 0.4 millimeters.

#### 5) Motion capture

We used the motion capture system to record the locations of 35 3D printed reflective markers that resembled a 4-millimeter reflective marker on top of a BioSemi active pin electrode (Figure 1F). We did not use actual BioSemi electrodes, which have wires that could prevent the cameras from seeing the markers. We placed 27 EEG electrode shaped markers to approximate the international 10-20 EEG cap layout and placed an additional eight EEG electrode shaped markers randomly to the cap to add asymmetry to improve tracking of the markers. We recorded 2-second takes of the positions of the 35 markers, three markers on the fiducials, and three face markers. Before transforming the locations to the head coordinate system, we identified and canceled movements of the head during data collection using the three face markers.

### B. Transformation to head coordinates

We developed a dedicated pipeline to convert the digitized electrode locations for each digitizing method to a format that could be imported to the common toolboxes for EEG analyses. Because EEGLAB and FieldTrip can easily read Zebris ElGuide’s output file (an SFP file), we created SFP files for all digitizations.

The head coordinate system in ElGuide defines the X-axis as the vector connecting the left preauricular to the right preauricular and the origin as the projection of the naison to the X-axis. Therefore, the Y-axis is the vector from the origin to the naison, and the Z-axis is the cross product of the X and Y unit vectors, which starts from the origin.

### C. Digitization reliability and validity

Variations in the digitized electrode locations could originate from random errors and systematic bias. The effects of random errors can be quantified as variability. Reliability is also inversely related to variability. Systematic bias can be quantified as the difference between measured locations and the ground truth locations. Validity is also inversely related to systematic bias.

#### 1) Digitization reliability

To assess the effects of random errors, we quantified digitization variability. We averaged the five digitized locations for each electrode to find the centroid. We then calculated the average Euclidean distances of the five digitized points to the centroid for each electrode and averaged those distances for all of the electrodes to quantify within-method variability. We identified and excluded outliers, single measurements that were beyond five standard deviations of the average variability for a digitizing method. If there were outliers, we recalculated the average digitization reliability with the updated dataset. Throughout the paper, we use “variability” to refer to “within-method variability.” Because reliability is inversely related to the variability, the most reliable method has the least variability.

#### 2) Digitization validity

To quantify the systematic bias of a digitizing method, we calculated the average Euclidean distance between the centroid for a digitizing method and the ground-truth centroid for the same electrode. We used the electrode centroids from the most reliable digitizing method as the ground-truth [43]. Then, we averaged the Euclidean distances for the 128 electrodes to obtain the magnitude of the systematic bias for each digitizing method. Because validity is inversely related to the systematic bias, the most valid method has the least systematic bias.

### D. Source estimation uncertainty

To generalize the possible effects of digitization reliability, we synthesized 500 sets of electrode locations with a Gaussian distribution using the variability average and standard deviation calculated for each digitizing method in II-C1. We excluded the motion capture method from the source estimation uncertainty analyses because we only recorded the locations of 35 EEG electrode shaped reflective markers instead of all 128 EEG electrode locations. We used a single representative 128-channel EEG dataset from a separate study for the source estimation analyses. We applied the Adaptive Mixture Independent Component Analysis (AMICA) to decompose EEG signals into independent components (ICs) [44], which has been reported to represent dipolar activities of different brain and non-brain sources [7].

We used EEGLAB’s DIPFIT toolbox version 2.3 to estimate a dipole equivalent for each IC and applied DIPFIT 500 times for each digitizing method. Each DIPFIT iteration used one of the 500 sets of synthesized electrode locations, the Montreal Neurological Institute (MNI) head model [45], and the ICs from the AMICA. The MNI head model is an averaged structural head model from 305 participants and provides 1mm × 1mm × 1mm resolution. To convert the mannequin head to be compatible with the MNI model, we warped the electrode locations to the MNI model using only the fiducials to preserve individual characteristics of the mannequin head. We used the dipoles produced with the electrode location centroids for the digitizing method with the highest reliability and identified the dipoles that described > 85% of the IC signal variance. We also excluded any dipole that was estimated to be outside of the brain volume for any of the DIPFIT results (500/method × five methods = 2500 DIPFIT results). In the end, 23 ICs remained.

#### 1) Spatial uncertainty

We fitted an enclosing ellipsoid with the minimum volume to each IC’s cluster of 500 dipoles [46] and quantified spatial uncertainty in terms of the volume and width of the ellipsoid. A larger ellipsoid volume indicated that a single dipole could reside within a larger volume, and thus, had greater volumetric uncertainty. A larger ellipsoid width indicated that a single dipole could have a larger shift in location. We calculated the ellipsoid’s width as the maximum distance that the IC’s dipoles could have from one another. We averaged the volumes and widths of all 23 ICs to quantify the spatial uncertainty for each digitizing method.

#### 2) Brodmann area accuracy

To identify Brodmann areas, we used a modified version of the `eeg_tal_lookup` function from EEGLAB’s Measure Projection Toolbox (MPT). This function looks for the anatomic structures and neighboring Brodmann areas in a 10-mm vicinity of each dipole, calculates the posterior probability that the dipole belongs to any of the identified Brodmann areas, and then assigns the dipole to the Brodmann area with the highest probability [47], [48]. We first set the “ground-truth” Brodmann area for each IC to be the Brodmann area identified with the IC’s dipole estimated using the centroid electrode locations from the most reliable digitizing method. We calculated Brodmann area accuracy as the percentage of the 500 Brodmann area assignments that matched the “ground-truth” Brodmann area.

We also analyzed Brodmann area accuracy using a template of electrode locations based on the MNI head model for source estimation [49]. Because the BioSemi 128-electrode cap is not based on the 10-10 electrode map, we non-rigidly warped the electrode locations from the most reliable digitizing method to match the MNI template. We compared the Brodmann area identified from the template to the “ground-truth” Brodmann area. Because the electrode locations for the template are exact, we could not calculate a percentage of assignments; thus, the template’s Brodmann area for each IC was either a hit or miss. However, we did calculate and compare the distance between the template’s dipole to the “ground truth” dipole with the distance between each digitizing method’s dipoles to the “ground truth” dipole. These distances indicated whether the dipoles were near the “ground truth” dipole.

### E. Statistical analysis

We used a one-way repeated measures analysis of variance (rANOVA) to compare the reliability and validity of the digitizing methods, the spatial uncertainty of the estimated dipoles, and the Brodmann area accuracy. For significant rANOVA’s, we performed Tukey-Kramer’s post-hoc analysis to determine which comparisons were significant. We also performed a one-sided Student t-test to identify if the Brodmann area accuracy of each digitizing method was different from the template. The level of significance for all statistics was α = 0.05.

Additionally, we fit a polynomial, using a step-wise linear model (MATLAB `stepwiselm` function), to describe spatial uncertainty as a function of digitization variability. We forced the y-intercept of the first-order polynomials and the y-intercept and y’-intercept of the higher-order polynomials to be zero. We set the y-intercepts to be zero for two reasons: 1) when we used the exact same electrode locations and performed DIPFIT 100 times, the maximum distance between source locations was on the order of 10^-4^ cm, and 2) the fit should not model the uncertainty values < 0 for positive digitization variability values. The step-wise linear model started with a zero order model and only added a higher-order polynomial term when necessary. The criterion for adding a higher-order polynomial term to the model was a statistically significant decrease of the sum of the squared error between the data points and the predicted values.

## III. Results

The variability for the five digitizing methods were visibly different, and electrodes located at the back of the head tended to have greater variability (Figure 2). The variability for the ultrasound method was generally largest compared to the other methods and could be as large as ~1.5 cm for electrodes at the back of the head. The variability for all electrodes digitized with the motion capture method was small, being no greater than 0.001 cm.

**Fig. 2.**
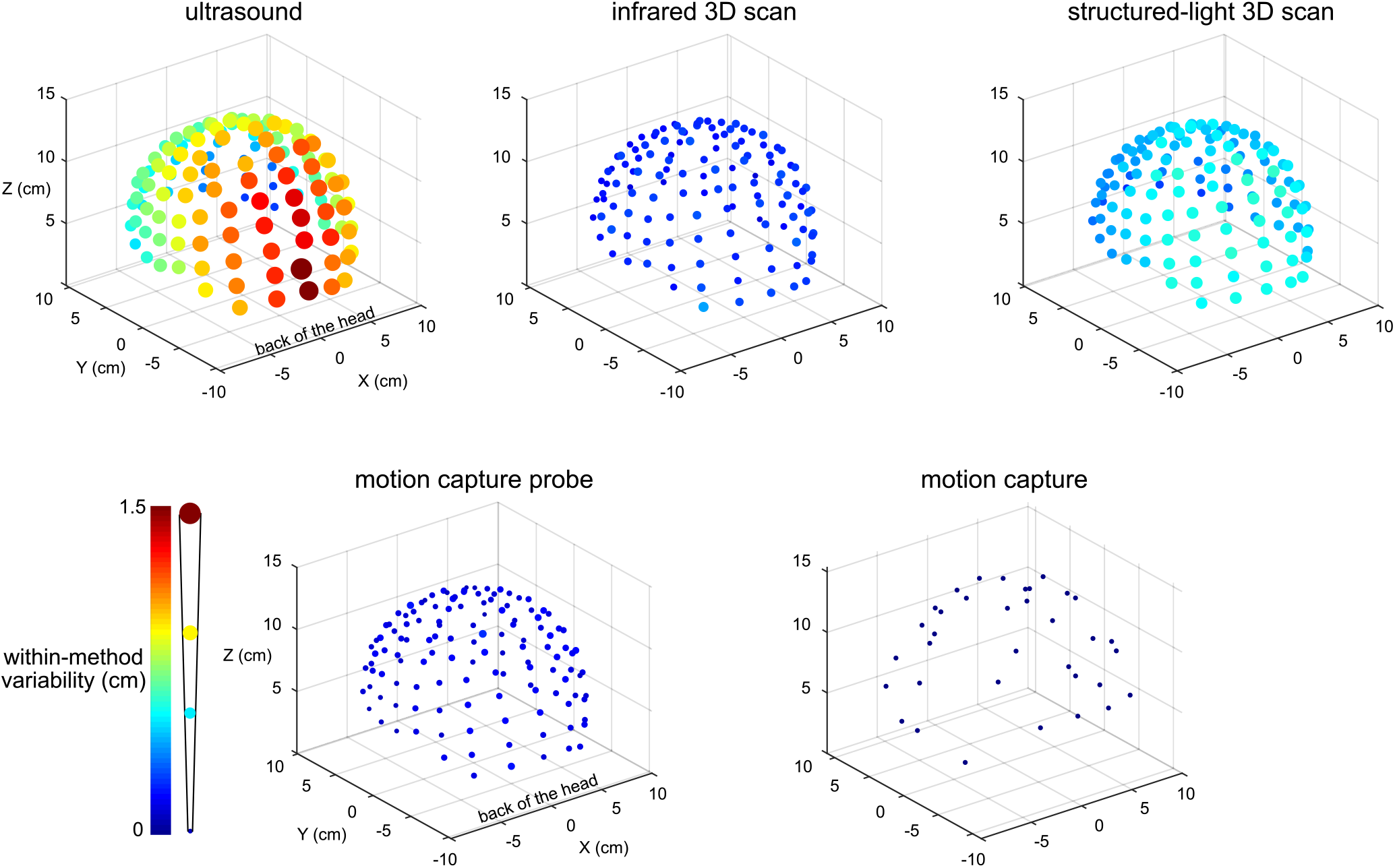
Visualization of the digitization reliability. Colored and scaled dots show the electrode location within-method variability for all 128 electrodes for the five digitizing methods. Ultrasound had the greatest variability and was the least reliable. The electrodes at the back of the head also tended to have the greatest variability. The motion capture method had the least variability and was the most reliable. The color bar and scale for the radii of the dots illustrate the magnitude of variability.

There was a range of reliability among the digitizing methods (Figure 3A). The motion capture digitizing method had the smallest variability of 0.001 ± 0.0003 cm (mean ± standard deviation) and hence, the greatest digitization reliability. The motion capture probe was the next most reliable method with an average variability of 0.147 ± 0.03 cm, followed by the infrared 3D scan (0.24 ± 0.05 cm), the structured-light scan (0.50 ± 0.09 cm), and the ultrasound digitization (0.86 ± 0.3 cm). The variability for the digitizing methods were significantly different (rANOVA F>1000, p<0.001), and the variability for each digitizing method was significantly different from all other digitizing methods (post-hoc Tukey-Kramer, p’s <0.001).

**Fig. 3.**
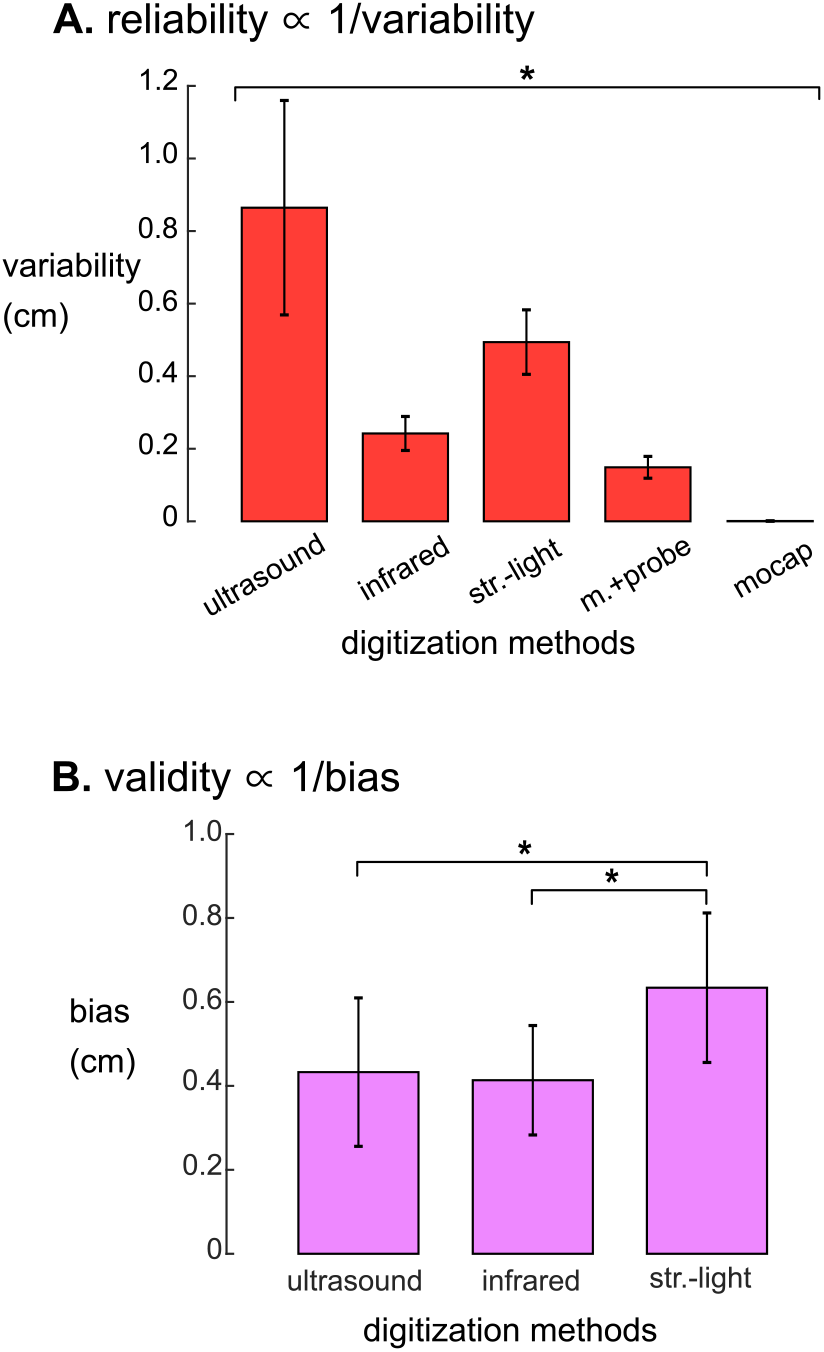
**A**: Reliability, quantified as the average variability, were significantly different for the five digitizing methods. The reliability of each digitizing method was significantly different from all other methods (* = Tukey-Kramer p’s <0.001 for all pair-wise comparisons). **B**: Validity, quantified as the average systematic bias showed that the structured-light 3D scan had the largest systematic bias compared to ultrasound and the infrared 3D scan. The motion capture probe method was assumed to be the ground truth and thus has no systematic bias and is not shown. * = Tukey-Kramer p’s <0.001. Error bars are the standard deviation. infrared = infrared 3D scan. str.-light = structured-light 3D scan. m.+probe = motion capture probe. mocap = motion capture.

The systematic biases, thus validities, of the digitizing methods were significantly different (rANOVA F>300, p<0.01, Figure 3B). The digitization validity of the structured-light 3D scan was the worst of the digitizing methods with a systematic bias of 0.63 ± 0.18 cm that was significantly larger than the other digitizing methods (post-hoc Tukey-Kramer, p’s <0.001). The digitization validity of the ultrasound and the infrared 3D scans were similar, with systematic biases of 0.43 ± 0.18 cm and 0.41 ± 0.13 cm, respectively.

Within a given digitizing method, dipoles generally showed similar spatial uncertainty while different digitizing methods generally showed differences in spatial uncertainty (Figure 4). Ellipsoid sizes for the motion capture probe, infrared 3D scan, structured-light 3D scan, and ultrasound digitization increased in order from the smallest to the largest, respectively. The enclosing ellipsoids of adjacent ICs also overlapped when the ellipsoid size was large, on the order of 1 cm^3^, such as for the ultrasound method.

**Fig. 4.**
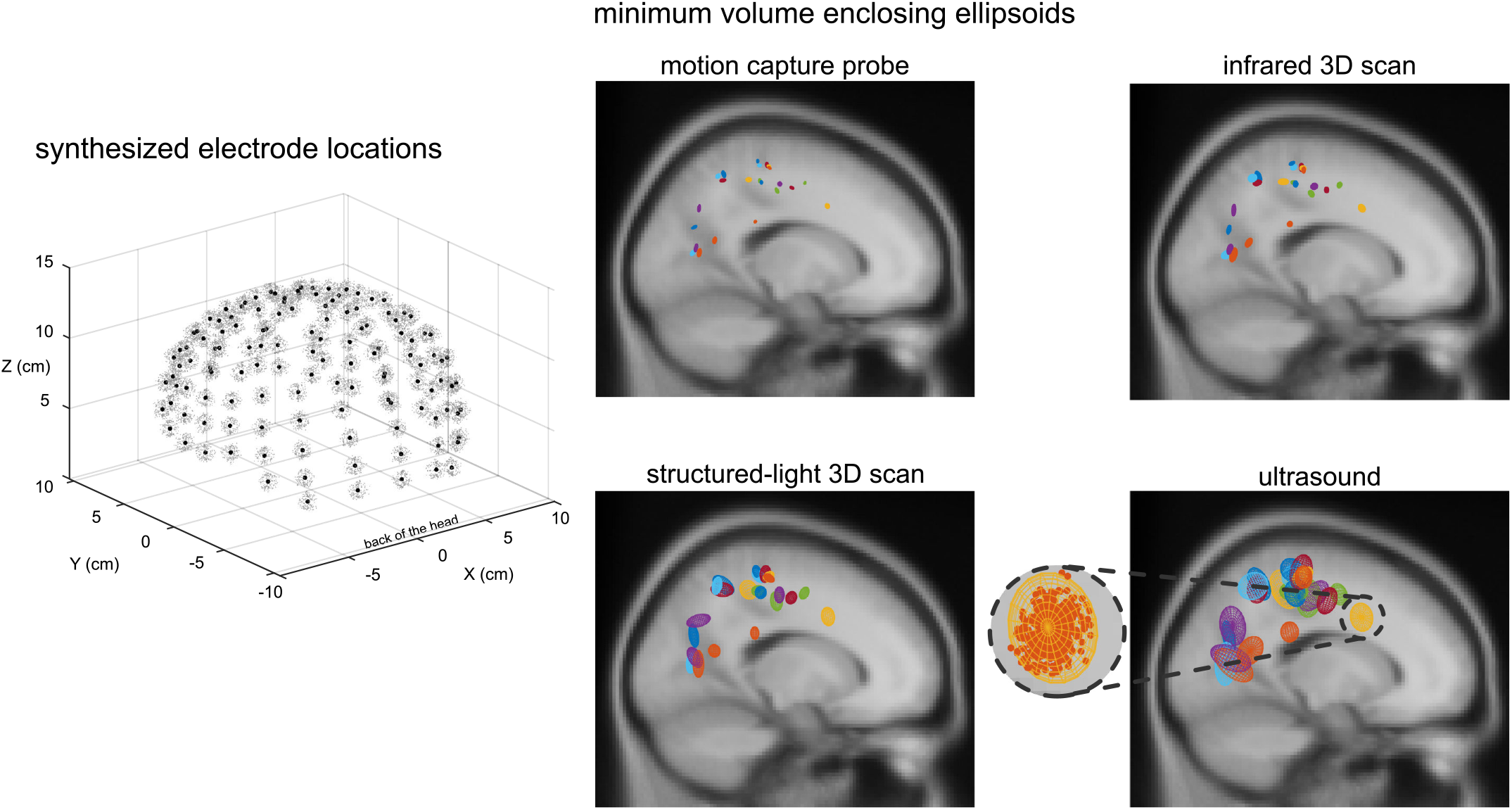
An example depiction of the synthesized electrode locations with a Gaussian distribution using the same averaged variability and standard deviation as the structured-light 3D scans, and the enclosing ellipsoids of the 500 dipoles for each independent component (IC) and digitizing method. Black dots = centroids of the electrode locations. Light gray dots = first 150 out of 500 synthesized electrode locations. Each color represents a different IC (23 ICs total). A close-up view of the ellipsoid fit for an Anterior Cingulate IC based on the reliability of the ultrasound digitizing method.

Ellipsoid volumes increased significantly with increasing digitization variability among the digitizing methods and had a cubic relationship (r^2^ = 1.00, Figure 5A). The motion capture probe and infrared 3D scan had the smallest uncertainty volumes (mean ± standard error) 0.007 ± 0.0007 cm^3^ & 0.029 ± 0.0027 cm^3^, respectively, whereas ultrasound had the largest uncertainty volume (1.37 ± 0.13 cm^3^). Structured-light 3D scan had an average uncertainty volume of 0.21 ± 0.014 cm^3^. The volumes of the enclosing ellipsoids showed a significant between-group difference (rANOVA, F>1000 p<0.001), and all uncertainty volume combinations of paired digitizing methods were significantly different (Tukey-Kramer post-hoc, p’s<0.001).

Ellipsoid widths also increased significantly with increasing digitization variability among the digitizing methods but had a linear relationship where the ellipsoid width was twice the size of the digitization variability (r^2^ = 1.00, Figure 5B). The average ellipsoid width was the smallest for the motion capture probe, (mean ± standard error) 0.34 ± 0.018 cm. The average ellipsoid widths for the two 3D scans were 0.53 ± 0.028 cm for the infrared 3D scan and 1.09 ± 0.051 cm for the structured-light 3D scan. The largest average ellipsoid width was for the ultrasound digitization, 1.90 ± 0.081 cm. The rANOVA for the widths of the enclosing ellipsoids showed a significant between-group difference (F>1000, p<0.001) and all combinations of paired digitizing methods had significantly different uncertainty widths (Tukey-Kramer post-hoc p’s<0.001).

**Fig. 5.**
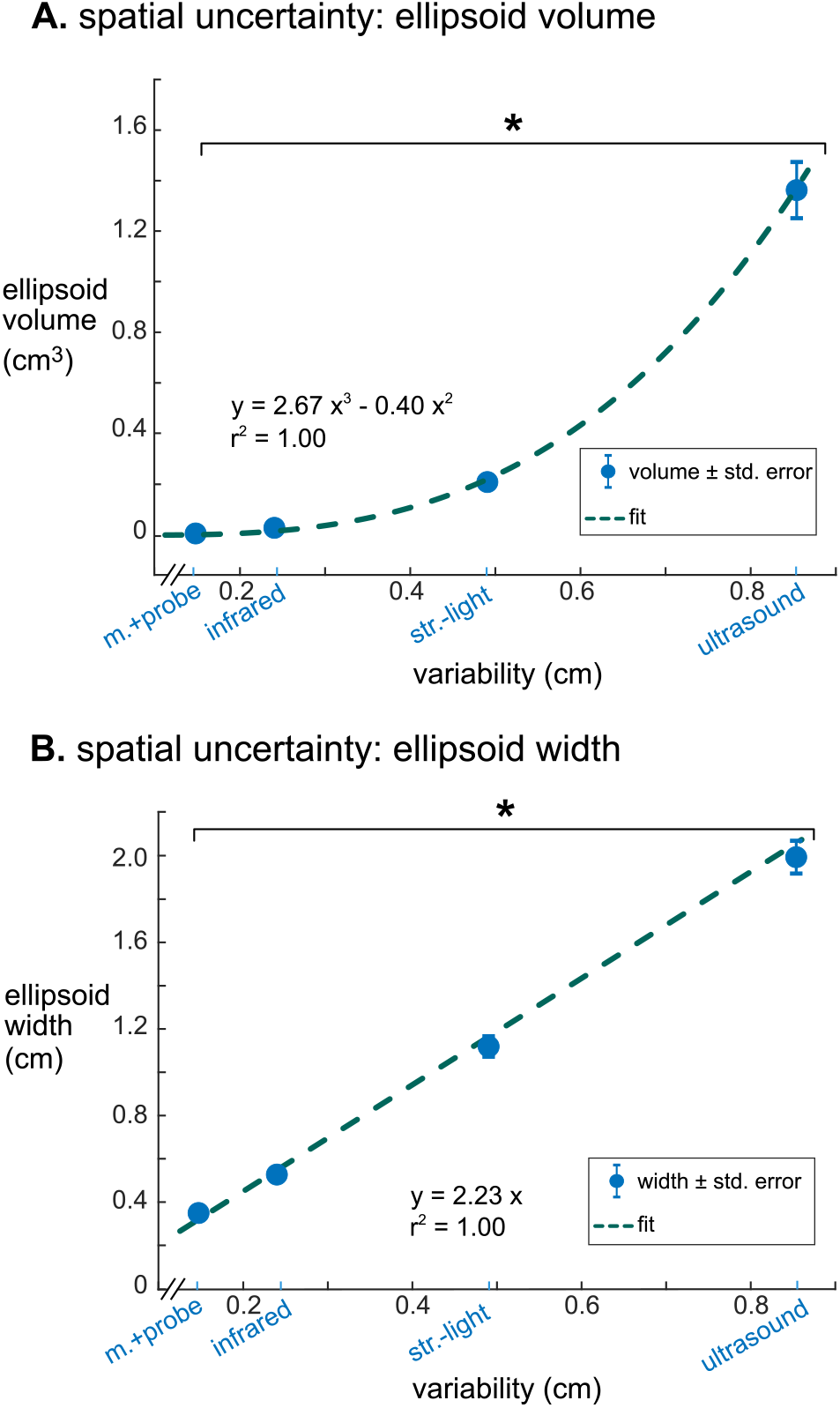
The relationships between digitization variability and dipole spatial uncertainty. **A**: Digitization variability and ellipsoid volume had a cubic relationship with an r^2^ of 1.00. **B**: Digitization variability and ellipsoid width had a linear relationship with an r^2^ of 1.00. Error bars are the standard error. * = Tukey-Kramer p’s <0.001 for all pair-wise comparisons. m.+probe = motion capture probe. infrared = infrared 3D scan. str.-light = structured-light 3D scan.

The Brodmann area accuracy among the digitizing methods could be extremely consistent within some ICs and could also be drastically different for other ICs (Figure 6 and in Supplement Figure S1). In general, the digitizing method with the highest reliability also had the highest Brodmann area accuracy within a given IC. For some ICs, all digitizing methods had > 98% Brodmann area accuracy. For other ICs, the Brodmann area accuracy decreased as reliability decreased. The most drastic example for this dataset was BA18 in Figure 6, where the Brodmann area accuracy was 86% with the motion capture probe method but dropped to 26% with the ultrasound method.

**Fig. 6.**
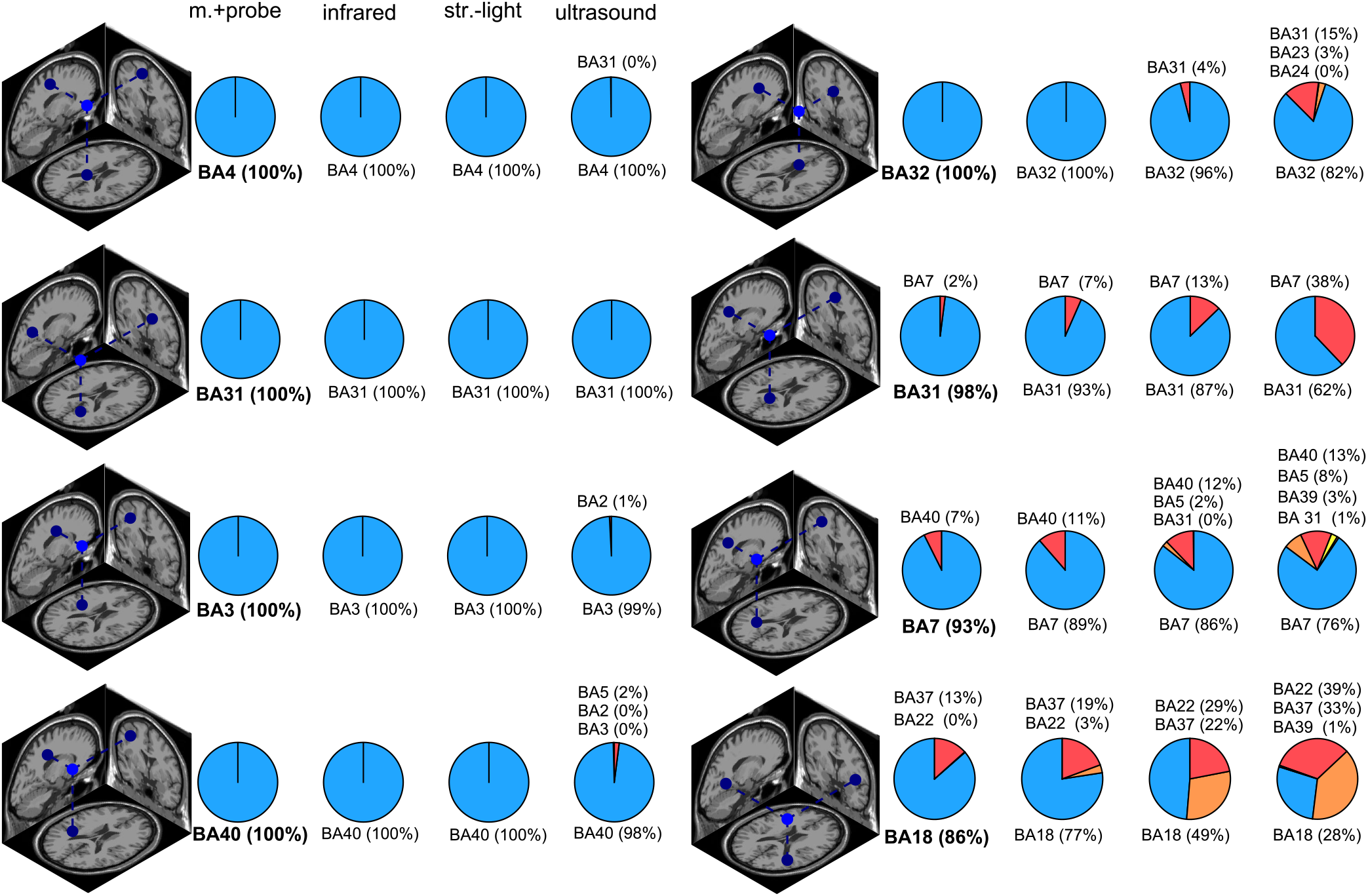
Brodmann area (BA) accuracy for a subset of ICs. The dipole depicts the “ground truth” dipole produced from the most reliable digitizing method, the motion capture probe method. The pie charts show the distribution of the Brodmann area assignments compared to the “ground truth” Brodmann area (shown in bold). ICs in the left column had consistent Brodmann area assignments regardless of digitizing method while the ICs in the right column had more varied Brodmann area assignments for the different digitizing methods. In general, less reliable digitizing methods led to less consistent Brodmann area assignments.

The Brodmann area accuracy for the digitizing methods and the template were significantly different (Figure 7). The motion capture probe had the highest Brodmann area accuracy, 93% ± 16 (mean ± standard deviation). The remaining digitizing methods in order of decreasing Brodmann area accuracy were the infrared 3D scan (91% ± 19%), the structured light 3D scan (87% ± 23%), and the ultrasound digitization (79% ± 25%). The rANOVA for the Brodmann area accuracy showed a significant between-group difference (F>306 p<0.001). Post-hoc Tukey-Kramer analysis showed significant pair-wise difference between all groups except the motion capture probe and infrared 3D scan. Using the MNI electrode template decreased the Brodmann area accuracy to 53% and was significantly different compared to any of the digitizing methods (p’s<0.001). The average distance of the dipoles of each digitizing method to the “ground-truth” dipole was less than 0.4 cm while the average distance of the template dipoles to the “ground-truth” dipole was ~ 1.4 cm.

**Fig. 7.**
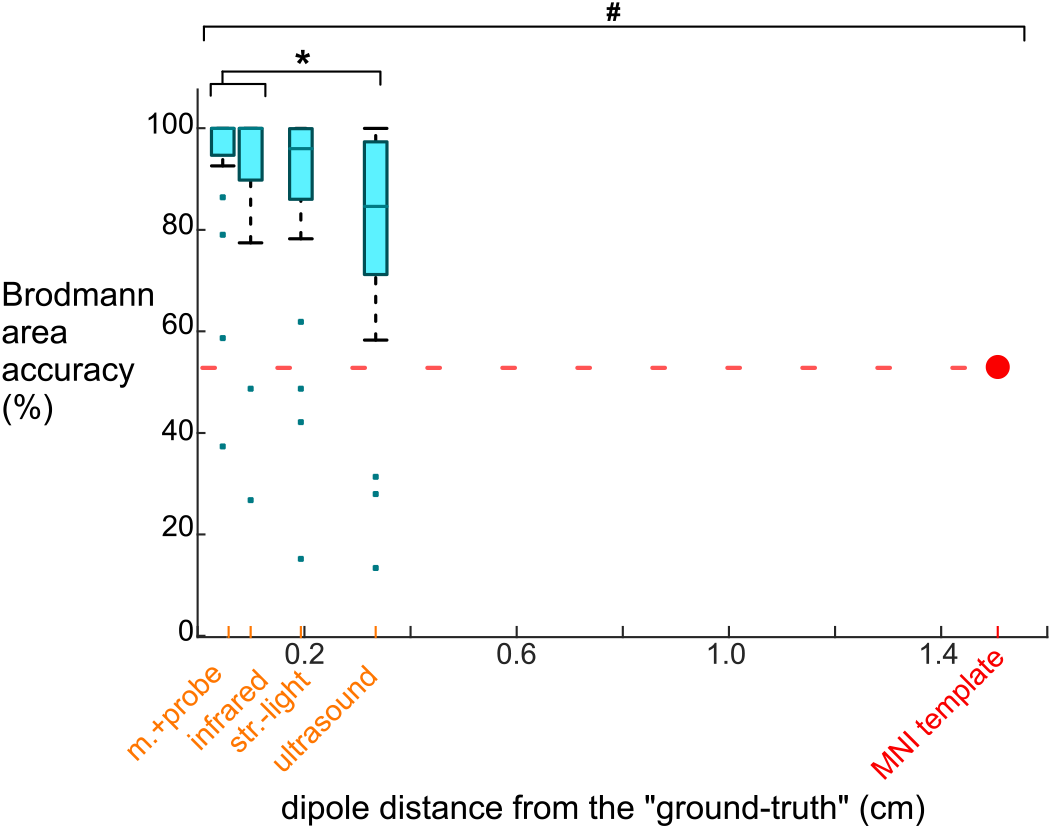
Brodmann area accuracy plotted versus the average dipole distance from the “ground truth” dipole when using different digitizing methods and the MNI template. Because larger distances between the dipoles and the “ground truth” likely would decrease Brodmann area accuracy, we plotted the methods on the x-axis at the method’s averaged dipole distance from the “ground truth” dipole. The box-whisker plot contains the Brodmann area accuracy averages for the 23 ICs. The Brodmann area accuracy average for an IC was the average of the percentage of the 500 iterations when the Brodmann area identified matched the “ground truth” Brodmann area for that IC. For the template, 53% of the Brodmann areas assigned for the 23 ICs using the template matched the “ground truth” Brodmann area. The Brodmann area accuracy was significantly different among the digitizing methods, except between the motion capture probe and infrared 3D scan (* Tukey-Kramer p’s <0.001). The template’s Brodmann area accuracy was significantly different than all digitizing methods (# Student’s t-test p’s <0.001). m.+probe = motion capture probe. infrared = infrared 3D scan. str.-light = structured-light 3D scan.

## IV. Discussion

We found that there was a range of reliability and validity among the digitizing methods and that less reliable digitizing methods translated to greater uncertainty in source estimation and poorer Brodmann area accuracy, assuming all other contributors to source estimation uncertainty were constant. Of the five digitizing methods (ultrasound, structured-light 3D scan, infrared 3D scan, motion capture probe, and motion capture), the most reliable digitizing method was the motion capture while ultrasound was the least reliable. The structured-light digitizing method had the greatest systematic bias and was thus the least valid method. We had hypothesized that less reliable digitizing methods would lead to greater source estimation uncertainty. In support of our hypothesis, digitizing methods with decreased reliability resulted in increased spatial uncertainty of the dipole locations and decreased Brodmann area accuracy. Surprisingly, any digitizing method led to an average Brodmann area accuracy of >80%. Using a template of electrode locations decreased Brodmann area accuracy to 53%. Overall, these results indicate that electrode digitization is crucial for accurate Brodmann area identification using source estimation and that more reliable digitizing methods are beneficial if the functional resolution for interpreting source estimation is more specific than Brodmann areas.

To help summarize the advantages of the different digitization systems, we created a table comparing the digitization reliability, dipole uncertainty, speed, affordability, and ease-of-use score, which are different factors that could influence which digitization a laboratory might choose to use (Table I). We estimated the digitizing speed as how much time each digitization required. The fastest digitizing method that required manual electrode marking was the motion capture probe method, which took 5 minutes to mark each electrode and 5 minutes to calibrate the system. The least expensive system was the infrared 3D scanner, which is likely to become even less expensive as cameras on smartphones become more advanced and could soon be used to obtain an accurate 3D scan for digitizing EEG electrodes. We also surveyed the operators to score each digitization on a scale of 1-5, with 1 being easy to use. While performing the actual 3D scan was perceived as being easy, marking the electrodes in MATLAB was not an easy task. The operators indicated that the motion capture was the easiest and that ultrasound was the most difficult method to use. To create a final ranking, we averaged the rankings for each factor (digitization reliability, dipole uncertainty, speed, affordability, and ease-of-use) to obtain a method score. Based on the method score, the best digitizing method was the motion capture. The next best method was tied between the motion capture probe and infrared 3D scan. The fourth best digitizing method was the structured-light 3D scan, and the worst digitizing method was the ultrasound method, which ranked poorly for all factors.

**TABLE I.** Rankings for each digitizing method based on factors related to performance, cost, and convenience. # is the rank of each method among all five methods and for the specified factor. The digitization reliability values and dipole uncertainty scalar width values were taken from our results. Speed was the approximate time a digitizing method required to obtain the file of electrode locations. The ease-of-use score was the average score operators provided in a survey with a score of 5 being the most difficult and 1 being the easiest method to do. The method score is the average rank of all factors for a given method and was defined as *score = Σ#/N*. Dipole uncertainty was not available for motion capture digitization. Mocap = motion capture. Infrared 3D = infrared 3D scan. Str.-Light 3D = structured-light 3D scan. * The probe price is for the OptiTrack digitizing probe. ** Motion capture cost was for an eight-camera system (optitrack flex13, $8000) and the optitrack motive software ($3000).

| method | digitization reliability |  | dipole uncertainty |  | speed |  | affordability |  | ease-of-use<br>(1 is easiest) | method score<br>(lower is better) |
| --- | --- | --- | --- | --- | --- | --- | --- | --- | --- | --- |
| | mean ± SD (cm) | # | mean ± SD (cm) | # | time (min) | # | cost (USD) | # | | |
| ultrasound | 0.86 ± 0.30 | 5 | 1.90 ± 0.39 | 4 | 30 | 5 | 15k | 5 | 4.2 | 4.6 |
| infrared<br>3D scan | 0.24 ± 0.05 | 3 | 0.53 ± 0.136 | 2 | 20 (10 for scan<br>+ 10 for marking<br>in software) | 3 | 1k | 1 | 2.5 | 2.3 |
| str.-light<br>3D scan | 0.50 ± 0.09 | 4 | 1.09 ± 0.24 | 3 | 25 (10 for scan<br>+ 15 for marking<br>in software) | 4 | 5k | 2 | 3.4 | 3.3 |
| mocap<br>probe | 0.15 ± 0.03 | 2 | 0.34 ± 0.08 | 1 | 10 (5 for digitizing<br>+ 5 for calibration) | 2 | 1k* (probe)<br>12k** (+mocap) | 4 | 2 | 2.2 |
| mocap | 0.001 ± 0.0003 | 1 | N/A | - | 5.1 (0.1 for digitizing<br>+ 5 for calibration) | 1 | 11k** | 3 | 1 | 1.5 |

For researchers interested in investing in having a digitizing system with the highest reliability, our results suggest that the motion capture method is currently the best option. In a previous study, Reis and Lochmann developed an active-electrode motion capture approach for an EEG system with 30 electrodes and reported small deviation of the digitized locations from the ground truth locations [31]. In addition to having sub-millimeter variability, the motion capture method only required 1-2 seconds to digitize, assuming that the markers were already placed on the EEG electrodes. However, tracking 64+ markers on an EEG cap will be challenging for most motion capture systems. In this study, we only tracked 35 markers on the EEG cap and may have been able to track more than 35 markers with our 22-camera motion capture system. Determining the maximum number of EEG electrodes that could be digitized using a motion capture approach could be beneficial and pursued in future work. Laboratories that already have a motion capture system and do not need to digitize more than 64 EEG electrodes could conveniently use the motion capture method, which would provide a cost-seffective, fast, and easy digitizing process. For laboratories that need to digitize 64+ electrodes and have a motion capture system already, the motion capture probe digitizing method would be the recommended option.

For researchers interested in investing in a cost-effective and convenient digitizing system, our results support recent efforts to use 3D scanners to digitize EEG electrodes. Both the structured-light and infrared 3D scanning methods were more reliable than digitizing with the ultrasound method. Furthermore, our reliability results for the two 3D scanners align well with a recent study that showed that an infrared 3D scan could automatically digitize electrode locations on three different EEG caps and achieve good reliability after additional post-processing [39]. Of the two 3D scanners we tested, the less expensive infrared 3D scanner was more reliable, had higher validity, and resulted in less dipole uncertainty, compared to the structured-light 3D scanner. Even though the structured-light 3D scanner provides more details from the mannequin head and cap, those details did not seem to be important for improving digitization reliability or validity. Additionally, the highly detailed structured-light 3D scans created large files and resulted sluggish refresh rates that made rotating and manipulating the 3D scan in MATLAB difficult. The infrared 3D scan, unlike the structured-light 3D scan, was in color, which was helpful for the operators to identify the EEG electrodes more easily on the computer screen. In the future, artificial intelligence approaches may be able to fully automate the digitizing process and use the additional topographic details from high resolution 3D scans. A continuous image-based digitizing method such as using a regular video recorded using a typical smartphone could also potentially be developed to digitize EEG electrode locations.

Compared to simulation studies, our experimental results demonstrated that source estimation uncertainty increased steeply with increasing EEG electrode variability. We showed that a digitizing method with an average variability of 1 cm could lead to a shift of a single dipole by more than 2 cm, which is > 20% of the head radius. There is just one simulation study that we know of that also showed a 2-fold increase in source uncertainty for every unit of digitization variability [22]. In that study, digitization variabilities were created using systematic rotations applied to every electrode location [22]. The majority of the simulation studies however, suggest that source uncertainty could only be as large as the digitization variability [19]–[21], [50]. In one of the mathematical studies, the theoretical lower bound of source estimation uncertainty was 0.1cm for 0.5cm shifts in EEG electrode location [21], which is 10x smaller than our experimental results. While simulation studies can be insightful, results should also be cross-validated with a conventional source estimation method (e.g. DIPFIT, LORETA or minimum norm) to determine whether simulation results are indicative of real-world source estimation uncertainty.

Because researchers often use Brodmann areas to describe the function of a source, we translated our results to be in terms of Brodmann area accuracy, which led to a few surprising revelations. The main revelation was that despite the range of digitization reliability, any of the digitizing methods we tested produced an average Brodmann area accuracy > 80%. As long as sources are only discussed according to Brodmann areas or larger cortical spatial regions, any current digitizing method can be used. The second revelation was that using the template electrode locations, instead of digitizing the electrodes, significantly decreased Brodmann area accuracy from > 80% to ~ 50%, which may be due to a ~ 1.5cm shift in dipoles locations (Figure 7). This shift may occur because the template removes information related to individual’s head shape. The third revelation was that for several sources, the same Brodmann area was almost always identified, regardless of the digitizing method used (left column in Figure 6). For other sources, less reliable digitizing methods led to more potential Brodmann area assignments (right column in Figure 6), but those different Brodmann areas may be functionally similar. Most likely, the proximity of a source to the boundary of a Brodmann area as well as the size of the Brodmann area contribute to Brodmann area accuracy. Ultimately, the accuracy of source estimation will depend on the target volumes of cortical regions of interest.

Limitations of this study were that we tested a subset of all digitizing methods, did not include digitization variability of the fiducials, and did not perform source estimation using other common algorithms. Even though we did not test many of the marketed digitizing systems, we replicated and tested the fundamental methods used by most of the marketed digitizing systems. One widely used EEG electrode digitizing method we did not test is an electromagnetic digitizing method (e.g. Polhemus Patriot or Fastrack system). Another study using similar digitization reliability analyses reported an average variability of 0.76cm for an electromagnetic digitization system [30], which is slightly better than the ultrasound digitizing method, with a variability of 0.86 ± 0.3cm. Here, the fiducials were nearly identical for every digitization, but in practice, marking the fiducials while the subject wears the cap can be challenging. Mismarking a fiducial can significantly shift the dipole location by 2 times the distance of the fiducial mismarking [51]. We excluded adding variations in fiducial locations to avoid confounding source estimation uncertainty due to digitization reliability and fiducial variability. Last, we did not use other different source estimation algorithms such as LORETA or beam-forming. Studies indicate that commonly used source estimation algorithms generally identify the similar source locations [52]–[54], which suggests that the choice of the source estimation algorithm used would probably not significantly alter our results.

Future efforts to improve source estimation, so that sources can be interpreted in terms of cortical spatial regions smaller than Brodmann areas, will involve more than just developing more reliable, convenient, and cost-effective digitizing methods to help reduce source estimation uncertainty. Even if a perfect digitizing method could be developed, there would still be uncertainty in source estimation as result of other factors such as improper head-model meshes and inaccurate electrical conductivity values [55], [56], which were assumed to have a constant contribution to the source estimation uncertainty in our analyses. Obtaining and using as much subject specific information, such as subject-specific MRI scans in addition to digitizing EEG electrode locations, should improve source estimation. EEGLAB’s Neuroelectromagnetic Forward Head Modeling Toolbox (NFT) could be used to warp the MNI head model to the digitized electrode locations to retain the individual’s head shape but is computationally expensive [57]. Using subject-specific MRIs instead of the MNI head model is also limited to groups with access to an MRI at an affordable cost per scan.

## V. CONCLUSION

In summary, there was a range of digitization reliabilities among the five digitizing methods tested (ultrasound, structured-light 3D scanning, infrared 3D scanning, motion capture with a digitizing probe, and motion capture with reflective markers), and less reliable digitization resulted in greater spatial uncertainty in source estimation and poorer Brodmann area accuracy. We found that the motion capture digitizing method was the most reliable while the ultrasound method was the least reliable. Interestingly, Brodmann area accuracy for a source only dropped from ~ 90% to ~ 80%, when using the most and least reliable digitizing methods, respectively. If source locations will be discussed in terms of Brodmann areas, any of the digitizing methods tested could provide accurate Brodmann area identification. Using a template of EEG electrode locations, however, decreased the Brodmann area accuracy to ~ 50%, suggesting that digitizing EEG electrode locations for source estimation results in more accurate Brodmann area identifications. Even though digitizing EEG electrodes is just one of the factors that affects source estimation, developing more reliable and accessible digitizing methods can help reduce source estimation uncertainty and may allow sources to be interpreted in terms of cortical regions more specific than Brodmann areas in the future.

---

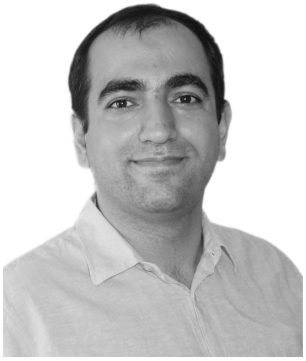

**Seyed Yahya Shirazi** was born in Tehran, Iran in 1988. He received his B.Sc. and M.Sc in biomedical engineering from Tehran Polytechnic in 2011 and 2014 respectively and is currently a Ph.D. student in mechanical engineering at the University of Central Florida (UCF).

Before joining UCF, his research focus was mainly on the postural stability of different clinical populations in perturbed conditions. His research interests include biomechanics and neuromuscular interactions of the human locomotion. Currently, he is a Graduate Assistant at the Biomechanics, Rehabilitation, and Interdisciplinary Neurosceince (BRaIN) laboratory at UCF and studies neuromechanical aspects of the human response to mechanical perturbations during locomotor tasks.

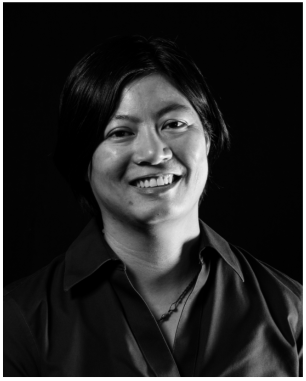

**Helen J. Huang** received her B.S. in materials science and engineering from the Massachusetts Institute of Technology in Cambridge, MA, USA in 2001 and her M.S. and Ph.D. in biomedical engineering from the University of Michigan, Ann Arbor, MI, USA in 2004 and 2009, respectively.

She was previously a Postdoctoral Research Associate and Fellow in the Integrative Physiology department at the University of Colorado, Boulder and an Assistant Research Scientist in the School of Kinesiology at the University of Michigan, Ann Arbor. She is currently an Assistant Professor in the Mechanical and Aerospace Engineering department at the University of Central Florida (UCF) where she also directs the UCF Biomechanics, Rehabilitation, and Interdisciplinary Neuroscience (BRaIN) Laboratory. Her research interests are neuromechanics, locomotion, motor adaptation, and gait rehabilitation.

## Notes

This work was supported by the National Institute on Aging of the National Institutes of Health, under award number R01AG054621. The content is solely the responsibility of the authors and does not necessarily represent the official views of the National Institutes of Health.

