## Supplementary Figures for "More Reliable EEG Electrode Digitizing Methods Can Reduce Source Estimation Uncertainty, But Current Methods Already Accurately Identify Brodmann Areas"

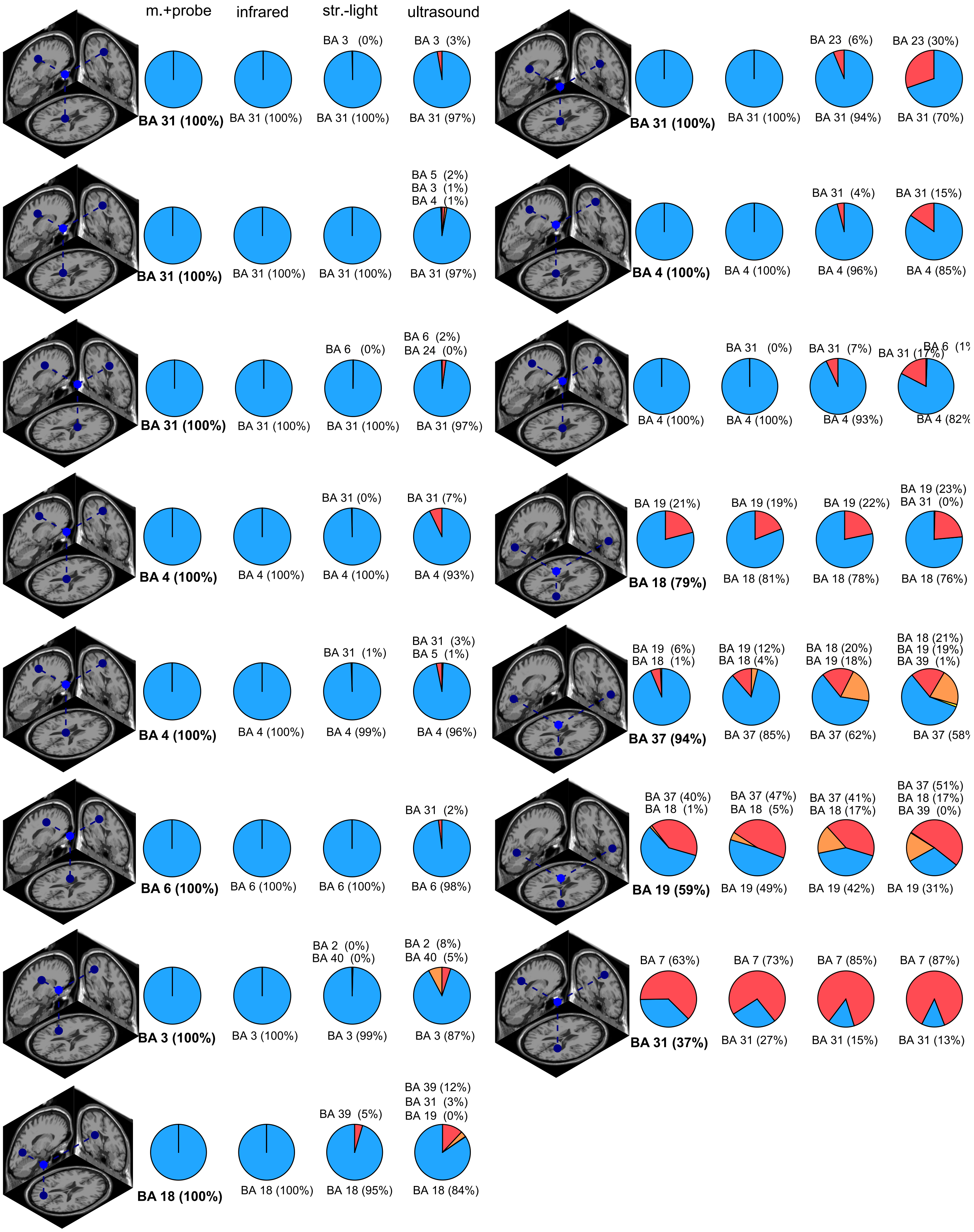


**Figure S1.** Brodmann area (BA) accuracy for the ICs not presented in the Figure 6.
